# Tracking Individual Differences in Perception by TMS-EEG Intrinsic Effective Connectivity

**DOI:** 10.1101/206797

**Authors:** Yuji Mizuno, Masahiro Kawasaki, Masanori Shimono, Carlo Miniussi, Yuka O Okazaki, Kenichi Ueno, Chisato Suzuki, Takeshi Asamizuya, Kang Cheng, Keiichi Kitajo

## Abstract

Non-invasive human electroencephalography (EEG) coupled with transcranial magnetic stimulation (TMS) is currently used to measure coarse stimulus-response relationships in brain physiology during behavior. However, with key modifications, the TMS-EEG technique holds even greater promise for monitoring fine-scale neural signatures of human behavior. Here, we demonstrate that a novel TMS-EEG co-registration technique can dynamically monitor individual human variation in perception based solely on EEG resting-state intrinsic effective connectivity probed by TMS-based phase resetting of ongoing activity. We used a bistable stimulus task, where the percept is perceived as either horizontal or vertical apparent motion, to record gamma band interhemispheric integration of information. Fine-grained inter-individual behavioral differences in horizontal motion bias could be measured by tracking resting-state gamma-band effective connectivity from right hMT+ to left hMT+. Thus, our method of triggering intrinsic resting-state effective connectivity in oscillatory dynamics can monitor individual differences in perception via the long-range integration of information. This technique will be useful for the manipulative dissection of individual-scale human cognition mediated by neural dynamics and may also expand neurofeedback approaches.

## Significance Statement

How individual differences in human behaviors emerge from brain networks is one of the most intriguing questions in human neuroscience. We demonstrate that a novel TMS-EEG technique can measure individual differences in effective connectivity of brain networks mediated by large-scale neural dynamics in the human brain. Using a bistable apparent motion stimulus, we assessed individual-level perceptual biases that are associated with the capacity for information integration across hemispheres. Inter-individual behavioral differences in motion bias could be tracked in intrinsic resting-state gamma-band effective connectivity from the right to left visual hemispheres. Our non-invasive perturbation method should be a useful tool for individual dissection of human brain dynamics which causally mediate information integration during perception and cognition.

## Introduction

The brain consists of functionally specialized modules localized to specific brain areas (Bullmore and Sporns, 2009). Long-range communication and integration of information between these modules is essential for brain functions including perception, action, and cognition. Synchrony in neural oscillations (Engel and Singer, 2001; Varela et al., 2001; Ward, 2003), which is a nonlinear dynamical phenomenon in the brain, is considered a critical mediator of communication and information integration between brain areas. For example, in face perception, large-scale neural network synchronization between parietal and occipitotemporal cortices is essential for information integration (Rodriguez et al., 1999). Likewise, perception of a bistable apparent-motion illusion is known to be associated with the modulation of coherence of gamma-band activity between left and right human middle temporal complex (hMT+) (Rose and Büchel, 2005).

In human neuroimaging, long-range communication and information integration can be measured by functional connectivity defined by the statistically-dependent activity between two neuronal groups. In contrast, effective connectivity refers to the measurement of directed connectivity, or a dynamic and causal chain of neural responses from one neuronal group to another, (e.g. see review Friston (Friston, 2011)). While functional connectivity is considered a good indicator of the general involvement of brain areas in behavior, effective connectivity is considered more meaningful to identify neuronal mechanisms in modular neural networks with directed information integration.

Recent technological developments based on concurrent transcranial magnetic stimulation (TMS) with electroencephalographic (EEG) recordings (TMS-EEG) (Ilmoniemi et al., 1997; Kitajo et al., 2015) have made it possible to assess effective connectivity in the human brain. Prior TMS-EEG studies have demonstrated the long-range propagation of TMS-evoked potentials and/or brain oscillations (Ilmoniemi et al., 1997; Komssi et al., 2002; Massimini et al., 2005; Rosanova et al., 2009; Thut et al., 2011; Bortoletto et al., 2015). Short rhythmic parietal TMS at an individual’s alpha frequency can entrain ongoing EEG oscillations in the stimulated cortical area (Thut et al., 2011). Moreover, single-pulse TMS can cause the transient phase resetting of neural oscillations at rest and the global propagation of phase resetting (Kawasaki et al., 2014). Thus, TMS-EEG is emerging as a versatile tool that can interfere with the phase of neural oscillations in a given area, via phase reset of neuronal activity, and concurrently track where and to what extent this perturbation propagates to.

Inter-individual variations are common in human behavior. For instance, inter-individual differences are observed in the ratio of the durations of horizontal and vertical apparent-motion perception (Chaudhuri and Glaser, 1991; Genç et al., 2011; Shimono et al., 2012). However, the physiological mechanisms responsible for individual variation in behavior remain unclear. One hypothesis is that information integration in neural oscillations differs across individuals for individual-level perception, However, there are no existing brain physiological methods to examine this conjecture and no prior reports demonstrate that intrinsic effective connectivity, the global propagation of phase resets from one region to another, can predict individual perception. Such a demonstration would show that TMS-EEG can manipulate and track individual-level network dynamics and behavior.

In this study, we report the development of a novel TMS-EEG co-registration methodology combining high-density EEG and TMS to monitor individual human variation in perception based solely on data from resting-state intrinsic effective connectivity in oscillatory dynamics. Using this method, we can for the first time directly measure interhemispheric effective connectivity by analyzing, on an individual basis, the degree of propagation of frequency-specific phase resetting between brain areas during rest and its relationship to individual behavioral differences.

## Materials and Methods

### Experimental Procedures

#### Participants

Twenty-one right-handed healthy participants with normal or corrected-to-normal visual acuity took part in the experiments. Three participants left the study after the fMRI experiment. The remaining eighteen participants (12 males) had an age range of 20-44 years (25.2 ± 6.0: mean ± s.d.). All participants provided written informed consent for each experiment and received monetary rewards after the experiments. The study was approved by the Ethical Committee of RIKEN. Figure 1 shows the overall experimental design.

#### fMRI (DDQ) experiment protocol

First, fMRI data were acquired while the participant performed a DDQ task. Visual stimuli were projected onto a translucent screen (visual angle: 29.12° × 36.65°). A gray fixation cross was also presented at the center of the screen (vertical line: 0.30°, horizontal line: 0.30°).

Figure 1B shows the task design for the fMRI (DDQ) experiment. In a DDQ task session, four stationary dots (dot diameter: 0.85°; luminance of filled gray dots: 1.05 cd/m^2^; luminance of black background: 0.04 cd/m^2^) and the gray fixation cross were initially presented in the central region (horizontal distance: 5.15°; vertical distance: 6.9°; aspect ratio: 1:1.34) of the screen for 30 s. Subsequently, DDQ stimuli were presented in the same central region of the screen for a total of 120 s. The DDQ stimuli were identical to those used by Rose and Büchel (2005). Briefly, two pairs of orthogonally located dots were alternately presented for 250 ms [i.e., stimulus onset asynchrony (SOA) = 250 ms]. This time was chosen because the effects of intentional control were relatively low during this interval (Shimono et al., 2011). The participant was instructed to report perception of either a horizontal motion or a vertical motion by pressing one of two keys. Each block consisted of five sessions. Three blocks, in a total of 15 sessions, were performed by all participants. A set of 3D T1 weighted high-resolution (1 × 1 × 1 mm^3^) anatomical images was acquired prior to the performance of Block 1.

All MRI images were acquired with a 4-Tesla whole-body MRI scanner (Agilent Technologies Inc., Santa Clara, CA, USA) using a transverse electromagnetic (TEM) volume coil as the transmitter (Takashima Seisakusho, Tokyo, Japan) and a 16-channel-array head coil as the receiver (Nova Medical Inc., Wilmington, MA, USA). During the scan, the participant was made to lie in a supine position on the patient table, with the head immobilized, and to wear earplugs to reduce the acoustic noise from the scanner. A four-shot echo-planar imaging (EPI) pulse sequence was used with the following parameters: 16 slices: slice thickness, 3 mm; repetition time (TR), 4465 ms; echo time (TE), 20 ms; flip angle (FA), 64.6°; field of view (FOV), 192 × 192 mm^2^, and matrix size, 128 × 128, resulting in an in-plane resolution of 1.5 × 1.5 mm^2^. The participant also wore an EEG cap with markers (vitamin E capsules) attached to the EEG electrode locations. In the following TMS-EEG experiment, the Brainsight TMS frameless coil navigation system (Rogue Research, Montreal, Quebec, Canada) was used to confirm that the positions of the markers did not deviate from the positions of the EEG electrodes. Respiration signals were recorded using a pressure sensor and cardiac signals using a pulse oximeter, and these physiological fluctuations were removed from the EPI images (Hu et al., 1995).

**Figure 1.**
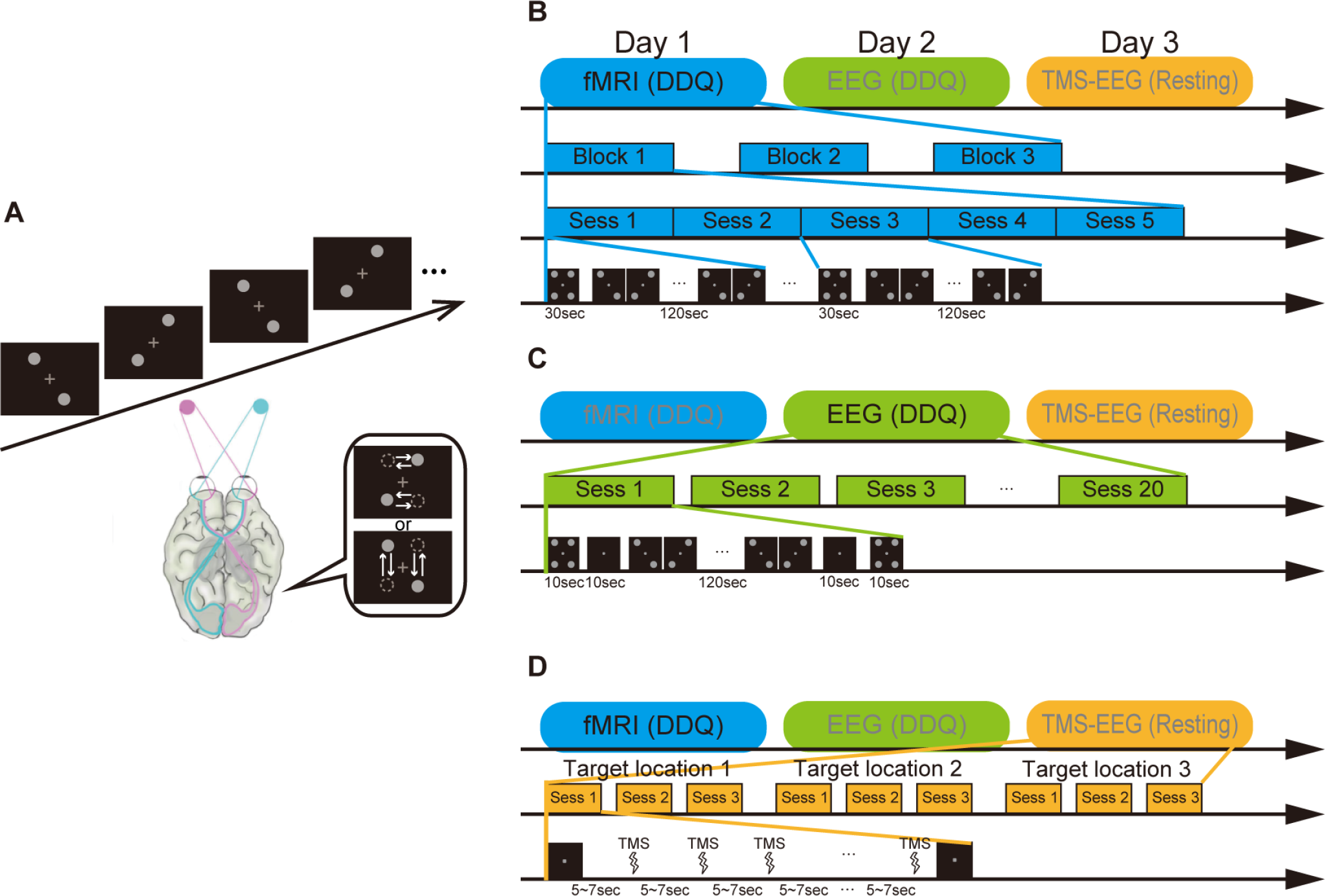
The Dynamical Dot Quartet (DDQ) task and experimental design. (**A**) DDQ stimuli consist of two alternately presented pairs of dots. Observers perceive either horizontal or vertical motion. Because visual information from each visual hemifield is processed in the contralateral hemisphere, information integration across the hemispheres is necessary for horizontal motion perception. The three experiments were performed on 3 separate days. (**B**) fMRI (DDQ) experiment. There were three blocks of the DDQ task, with each block consisting of five sessions. In each session, first, four dots and a fixation cross were presented for 30 s. Subsequently, the DDQ stimuli were presented for a total of 120 s. The DDQ stimuli alternated every 250 ms. (**C**) EEG (DDQ) experiment. EEG (DDQ) was performed for 20 sessions, 10 neutral and 10 biased sessions. The order of neutral and biased sessions was randomized and counter-balanced. Each session consisted of five parts. First, four dots and a fixation cross were presented for 10 s. Second, the fixation cross alone was presented for 10 s. Third, the DDQ stimuli were presented for 120 s. Then, the fixation cross was presented again for 10 s. Finally, four dots and the fixation cross were presented for 10 s. The DDQ stimuli alternated every 250 ms. (**D**) TMS-EEG (resting) experiment. TMS-EEG (resting) was performed for three consecutive sessions for each of three target locations (right hMT+, left hMT+, and sham). The order of the TMS target locations was randomly determined. In each session, participants were given 30 TMS pulses at intervals of 5 s to 7 s.

#### EEG (DDQ) experiment protocol

EEG was recorded during the DDQ task, where the DDQ stimuli and fixation cross were presented centrally on the black background of a CRT monitor (CPD-E220, Sony, Japan; refresh rate: 60 Hz, resolution: 1024 × 768, black background: 0.1 cd/m^2^). A chin rest was used to maintain head position at a distance of 100 cm from the monitor. In a task session (Fig. 1C), four dots (vertical distance: 6.9°, horizontal distance: 5.15°, aspect ratio: 1:1.34, diameter: 0.85°, filled gray circles: 19.8 cd/m^2^) and a gray fixation cross (19.8 cd/m^2^) were initially presented at the center of the monitor for 10 s. The fixation cross alone was presented at the center of the screen for 10 s. Then, the DDQ stimuli were presented for a total of 120 s, alternating every 250 ms. After the DDQ presentation, the fixation cross was presented for 10 s and then the four dots plus fixation cross were presented for 10 s. Each session was assigned as either ‘neutral’ or ‘biased’. In the neutral condition, participants were instructed to look at the DDQ stimuli passively without any intention and report their perception. In the biased condition, participants were asked to bias their perception towards horizontal motion for as long as possible. Each participant performed ten neutral and ten biased sessions in a random and counter-balanced fashion. In both conditions, participants reported their perceptual interpretation (i.e., horizontal motion or vertical motion) by depressing one of two keys. In this study, the analyses were focused on data from the neutral-condition sessions.

High-density EEG signals (63-channel) and horizontal and vertical EOG signals were recorded using a TMS-compatible EEG amplifier (BrainAmp MR+, Brain Products, Gilching, Germany) with a sampling rate of 1000 Hz with an on-line band-pass of DC-1000 Hz, and Waveguard caps with Ag/AgCl sintered electrodes (ANT Neuro, Enschede, The Netherlands) according to the international 10-10 system. AFz was used as the ground electrode, the left mastoid was used as a recording reference, and EEG data were re-referenced offline to the average of the left and right mastoid electrodes.

#### TMS-EEG (resting) experiment protocol

The TMS-EEG experiment was conducted during the resting (task-free) state. The EEG setup was the same as in the EEG (DDQ) experiment. Participants sat on a chair with their eyes open. They wore earplugs to minimize the effects of the TMS clicking sound. A fixation cross was presented centrally on the black background of a CRT monitor (CPD-E220, Sony, Tokyo, Japan, refresh rate: 60 Hz, resolution: 1024 × 768). Single-pulse TMS pulses (Magstim Rapid, The Magstim Company, Whitland, U.K.) were delivered to the left hMT+ and the right hMT+. The Brainsight frameless coil navigation system was used to guide the TMS coil to left and right hMT+ using the individual positions of peak hMT+ activation determined for each participant in the fMRI experiment. In the real TMS condition, the TMS coil handle was pointed upward, as in previous studies (Ruzzoli et al., 2010; Guzman-Lopez et al., 2011). In the sham condition, the TMS coil was positioned 10 cm above the vertex of the scalp and the TMS coil handle was pointed backwards. In all conditions, TMS pulses were delivered at intervals from 5 to 7 s with an intensity of 90% of the active motor threshold. To determine the active motor threshold, single-pulse TMS was applied over the left motor cortex. The initial stimulator output was set to 70% of the maximum stimulator output, and the pulse interval was 2 or 3 s. The active motor threshold for each individual was determined as the intensity to evoke a minimally perceptible twitch of the right index finger muscles in two consecutive trials while the participant performed weak isometric index finger contraction. Each session was composed of 30 TMS pulses (Fig. 1D). Participants were given 90 TMS pulses (30 pulses × 3 sessions = 90 pulses) for each target location (right hMT+, left hMT+, and sham). The order of the TMS target locations was random and counter-balanced. Data from three participants were removed from the group data analyses: two participants had excessive muscle artifacts in their TMS-EEG (resting) data (the number of survived epochs after removing artifact-contaminated epochs was less than 10), and one participant had a very low perceptual alternation rate in the EEG (DDQ) experiment (mean perceptual intervals for horizontal and vertical motion were larger than 50 s). In addition, data from three participants were removed from TMS-EEG data analyses because one participant did not show reduced TMS-evoked decay artifacts which lasted longer than 300 ms after an independent component analysis (ICA)-based artifact removal procedure and two participants did not indicate a *ZPLF* peak based on the criterion (*ZPLF* > 2).

#### fMRI (DDQ) data analysis

Bilateral hMT+ were localized in each participant to determine the optimal location for the TMS coil and nearby EEG electrodes. All fMRI data were processed and analyzed with SPM8 (Wellcome Institute of Cognitive Neurology, London, UK, http://www.fil.ion.ucl.ac.uk/spm/software/spm8) on an individual basis, including slice timing correction, realignment to correct for motion within the individual participant, coregistration to the initial scan, segmentation, and spatial smoothing with a Gaussian kernel of 8 mm full width at half maximum. Individual analyses were based on the general linear model, and task stimulus periods were convolved with the canonical two-gamma hemodynamic response function. The estimated head motion parameters were added as additional regressors. The statistical test was done between the baseline period (four dots and a fixation cross) and the DDQ period. The threshold for reporting significant activations was severally controlled to *p* < 0.001, corrected for multiple comparisons (FWE: Family-wise error rate corrected) based on random field theory. TMS was delivered to the peak of the defined activated areas.

### EEG (DDQ) data analysis

#### Behavioral analysis

Horizontal motion perception and vertical motion perception were extracted from the key-press data. Then, the total durations of perceived horizontal motion and perceived vertical motion were calculated. Perceptual bias was calculated as:

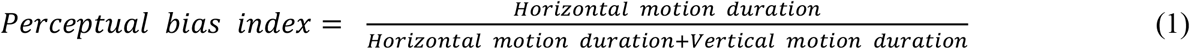

Higher values on the perceptual bias index indicate stronger horizontal bias in DDQ perception. This index was used to evaluate individual perceptual bias.

#### EEG data analysis

First, a 0.1 Hz to 100 Hz band-pass filter and a 47 Hz to 53 Hz notch filter were applied across all data. Next, the horizontal perception periods and vertical perception periods were extracted separately in 4000 ms after an indicated perceptual alternation. Epochs were obtained from intervals in which either horizontal or vertical motion perception was sustained for longer than 4000 ms. Then, an ICA-based automatic artifact removal procedure (ADJUST) was applied to the extracted epochs to remove generic artifacts, eye movements, and blink-related artifacts (Mognon et al., 2010). After the ICA-based procedure, epochs that contained signal amplitudes over 150 μV or below −150 μV were rejected to remove the remaining muscle artifacts. Then, the EEG data were transformed to current source density (CSD) data (Perrin et al., 1989, 1990)using the CSD Toolbox (Kayser and Tenke, 2006) The number of epochs in horizontal and vertical motion condition were equalized to the minimal number of epochs within each participant (mean: 32.9 ± 12.1: mean ± s.d.). Instantaneous phase was estimated using the Gabor wavelet from 4 Hz to 45 Hz at 1 Hz steps (Nco: 8). The central 2000 ms of all epochs was used after calculating phase to remove the edge effect of the wavelet transform. The phase-locking value (*PLV*) was computed as a measure of large-scale neural synchrony (Lachaux et al., 1999; Rodriguez et al., 1999). *PLV* was calculated as follows:

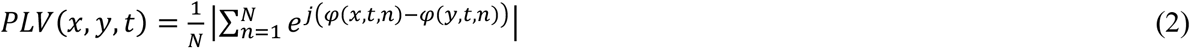

where, *x* and *y* are the indices of the electrode, *t* is time, *f* is frequency, *N* is the total number of epochs, and *φ* is the instantaneous phase of the CSD-transformed EEG. The *PLV* was estimated for each participant and each period. The *PLV* is strongly biased by the number of epochs, which in turn differed across participants. To eliminate this effect, the *PLV* was transformed to Rayleigh’s *z* value (*ZPLV*) using *ZPLV* = *N* × *PLV*^2^, where *N* is the total number of epochs. The temporally averaged *ZPLV* for horizontal motion perception and vertical motion perception were statistically compared with the cluster-based permutation test based on clustering of adjacent frequency points and electrode pairs proposed by Maris et al. (Maris and Oostenveld, 2007; Maris et al., 2007) (one-tailed cluster significance threshold: *P* < 0.05, one-tailed *t*-test, cluster-forming threshold: *P* < 0.05, *n* = 15, 10,000 permutations). Here we assume that all interhemispheric electrode pairs from EOIs are located adjacent each other. This method was applied to identify significant differences in *ZPLV* clusters between horizontal and vertical perception periods across electrode pairs and frequencies over all participants.

#### TMS-EEG (resting) data analysis

TMS-evoked exponentially decaying EEG artifacts were removed on a single-trial basis by modifying the ICA-based method (Herring et al., 2015). First, data epochs were extracted from −3000 ms to 3000 ms around the TMS pulses. Second, linear interpolation was performed from 0 ms to 4 ms, 6 ms, 8 ms, 10, or 12 ms after TMS onset (pre-linear interpolation). Later, one of the five interpolation intervals was selected. Third, ICA was applied to the linearly interpolated single-epoch data using the FieldTrip toolbox (http://fieldtrip.fcdonders.nl/) (Oostenveld et al., 2011). Then, artifact independent components (ICs) were removed as follows. First, the maximum value was detected between 0 ms and 100 ms in each IC. The maximum IC values were transformed to *z* scores with the mean and s.d. calculated across the all maximum IC values. ICA components were removed if the *z* scores were greater than 1.65. To eliminate remaining EEG artifacts, linear interpolation was performed from 0 ms to 12 ms (post-linear interpolation).

Then, a 0.01 Hz to 100 Hz band-pass filter and a 48 Hz to 54 Hz notch filter were applied across all data. The −1500 ms to −1000 ms period was used as a baseline period. Epochs were rejected if they contained signal amplitudes over 150 μV or below −150 μV within the 0 ms to 500 ms period after TMS and during the baseline period. Then, the CSD was computed (Perrin et al., 1989, 1990; Kayser and Tenke, 2006). The number of epochs used for each condition was the unified minimal number of epochs (56.5±10.2: mean ± s.d. *n* = 12) within each participant. Instantaneous phase was estimated using a Gabor wavelet from 4 Hz to 45 Hz at 1 Hz steps (Nco: 3). After calculating phase, only the −2000 ms to 2000 ms period of the 6000 ms epoch was used to eliminate the edge effects from the wavelet transform. Next, the phase-locking factor (*PLF*) (Tallon-Baudry et al., 1996) was computed as follows:

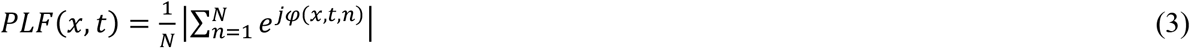

where *x* is the index of the electrode, *t* is time, *f* is frequency, *N* is the total number of epochs, and *φ* is the instantaneous phase of the CSD-transformed EEG. The *PLF* was transformed to Rayleigh’s *z* value (*ZPLF*) using *ZPLF* = *N* × PLF^2^ to eliminate the effect of the number of epochs, where *N* is the total number of epochs. Auditory evoked potentials caused by the TMS clicking sound were generated approximately 80 ms after the TMS pulses (Nikouline et al., 1999; Morishima et al., 2009). The *ZPLF* time course was obtained by averaging *ZPLF* across the electrodes in the EOIs (P6, P8, and PO8 electrodes for the right hMT+ and P5, P7, and PO7 electrodes for the left hMT+) and frequencies corresponding to the significant cluster obtained in the EEG (DDQ) experiment separately for each hemisphere.

The *ZPLF* time courses for the right hMT+ and left hMT+ in the real TMS and sham conditions were statistically compared with the cluster-based permutation test based on clustering of adjacent time points (Maris and Oostenveld, 2007; Maris et al., 2007) (one-tailed cluster significance threshold: *P* < 0.05, one-tailed *t*-test, cluster-forming threshold: *P* < 0.05, *n* = 12, 10,000 permutations). For further analyses, the *ZPLF* from the sham condition was subtracted from the *ZPLF* from the real TMS sessions to remove the effects of the TMS clicking sound. Brainstorm was used for topographic mapping of *ZPLF*s (Tadel et al., 2011).

Then, the contralateral *ZPLF* peak was normalized according to the following equation:

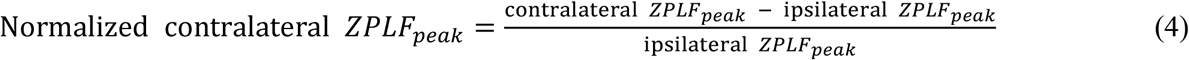

where ipsilateral *ZPLF*_peak_ is the peak *ZPLF* ipsilateral to TMS, and contralateral *ZPLF*_peak_ is the peak *ZPLF* contralateral to TMS.

The ipsilateral *ZPLF*_peak_ was defined as follows. First, the peak *ZPLF* was detected from 0 ms to 100 ms in each pre-linear interpolation (4 ms, 6 ms, 8 ms, 10 ms, or 12 ms). Second, the pre-linear interpolation intervals that showed peak *ZPLF* over 50 or below 2 were rejected assuming that EEG signals were contaminated by artifacts. Finally, the pre-linear interpolation interval that showed the shortest latency in peak time among the remaining datasets was selected. The contralateral *ZPLF*_peak_ was defined as follows. First, the peak *ZPLF* was detected in a time period from 12 ms to 150 ms and at least 15 ms after ipsilateral *ZPLF*_peak_ time. Next, the pre-linear interpolation interval that showed the shortest latency in peak time among the five datasets was selected.

Normalized contralateral *ZPLF*_peak_ values were calculated using the electrode pairs and frequency bands that showed significant clustering in the EEG (DDQ) experiment. Normalized contralateral *ZPLF*_peak_ in right and left TMS conditions were statistically compared using a Wilcoxon signed-rank test (*p* < 0.05, two-tailed). Finally, we analyzed the relationships between the perceptual bias index and normalized contralateral *ZPLF*_peak_ using the Spearman’s correlation coefficient (*p* < 0.05 one-tailed, Bonferroni-corrected for two TMS-targeted sites).

We also estimated *ZPLV* for −2000 ms to −1500 ms inter-stimulus resting intervals to assess intrinsic functional connectivity. *ZPLVs* were averaged across the electrode pairs and frequency bands that showed significant clustering in the EEG (DDQ) experiment.

## Results

### fMRI (DDQ) Experiment for Task-based Functional Localization

We performed a functional magnetic resonance imaging (fMRI) experiment using a dynamical dot quartet (DDQ) stimulus [fMRI (DDQ) experiment] to localize individual hMT+ to select electrodes of interest for subsequent EEG experiments and individualized TMS coil navigation. The DDQ task consists of two pairs of diagonally located dots, presented in an alternating sequence (Chaudhuri and Glaser, 1991; Rose and Büchel, 2005; Genç et al., 2011; Shimono et al., 2012) (Fig. 1A). Observers perceive either horizontal motion, where the dots are perceived as alternately passing across opposite visual hemifields, or vertical motion, where the dots are perceived as moving within the visual hemifield, and the direction of motion perception alternates spontaneously. Gamma-band synchrony between EEG electrodes over the left and right lateral parietal areas, encompassing hMT+ (P7-P8), is further increased during the perception of horizontal compared to vertical motion(Rose and Büchel, 2005) (i.e., inter- vs. intra-hemispheric integration) using 34-ch EEG recordings. However, for our purpose, it was necessary to conduct an individual-level task-based functional localizer to detect electrodes of interest (EOI) for 63-ch high-density EEG and individualized TMS coil navigation.

Participants were shown DDQ stimuli with the frames alternating every 250 ms. We identified individual left and right hMT+ locations by analyzing peak activations during apparent motion perception. Supplementary table 1 shows the group-averaged left and right locations of peak activations in MNI coordinates. Corresponding t-values (FWE-corrected) for activations were 10.7 ± 3.6 for Left MT+ and 10.9 ± 4.6 (mean ± s.d.) for Right hMT+, respectively (*n*= 15). Bilateral hMT+ showed significant activation when participants perceived apparent motion in the DDQ. The locations were consistent with previous fMRI studies (Dumoulin et al., 2000; Sterzer et al., 2003). Left hMT+ was located near the P7, P5, and PO7 electrodes, and Right hMT+ was located near the P8, P6, and PO8 electrodes. The locations were also consistent with estimated cortical projections of EEG electrodes (Koessler et al., 2009). In subsequent experiments, we applied TMS to locations on the scalp closest to the individual left and right hMT+. For further EEG analyses for EEG and EEG-TMS experiments, these electrodes were used as the electrodes of interest (EOIs).

### EEG (DDQ) experiment to select frequency bands and electrode pairs

We performed an EEG experiment using DDQ to evaluate changes in interhemispheric functional connectivity in oscillatory dynamics between left and right hMT+ in horizontal motion perception and vertical motion perception. We also evaluated the degree of individual perceptual bias via key-presses (left vs. right response for horizontal vs. vertical motion perception, respectively). We assessed the inter-hemispheric phase synchronization of current source density (CSD)-transformed EEG activity between EOIs over the right and left MT+ areas during DDQ perception. Figure 2 shows interhemispheric phase synchrony of EEG activity in horizontal motion perception and vertical motion perception. The phase synchrony between left and right hemispheres was estimated as the Rayleigh *z* value for the phase-locking value (Lachaux et al., 1999; Rodriguez et al., 1999) (*ZPLV*) and temporally averaged for each horizontal and vertical motion perception. The interhemispheric phase synchrony indicated a significant group difference between the perception of horizontal and vertical motion, assessed by a cluster-based permutation test (Maris and Oostenveld, 2007; Maris et al., 2007) (*P* = 0.020: one-tailed cluster test, *n* = 15, 10,000 permutations, cluster-forming threshold *P* < 0.050: one-tailed *t*-test). The corresponding electrode pairs and frequency bands in the cluster were as follows: P7-P8: 28 Hz to 45 Hz, P7-P6: 29 Hz to 36 Hz, P7-PO8: 45 Hz, PO7-P8: 32 Hz to 34 Hz, P5-P6: 33 Hz to 36 Hz, PO7-P6: 36 Hz to 38 Hz, and PO7-PO8: 30Hz, 31Hz, 36 Hz to 45 Hz. In Figure 2, *ZPLV* was averaged across the interhemispheric electrode pairs around hMT+ and across participants. Group-averaged perceptual intervals for horizontal motion perception and vertical motion perception were 1,012 ± 613 ms and 1,095 ± 629 ms (mean ± s.d. *n* = 15), respectively. The perceptual bias index, which indicates the individual degree of horizontal motion bias and takes a value of 0.5 for no bias, showed individual differences ranging from 0.35 to 0.58 (0.49 ± 0.06: mean ± s.d. *n* = 15).

**Figure 2.**
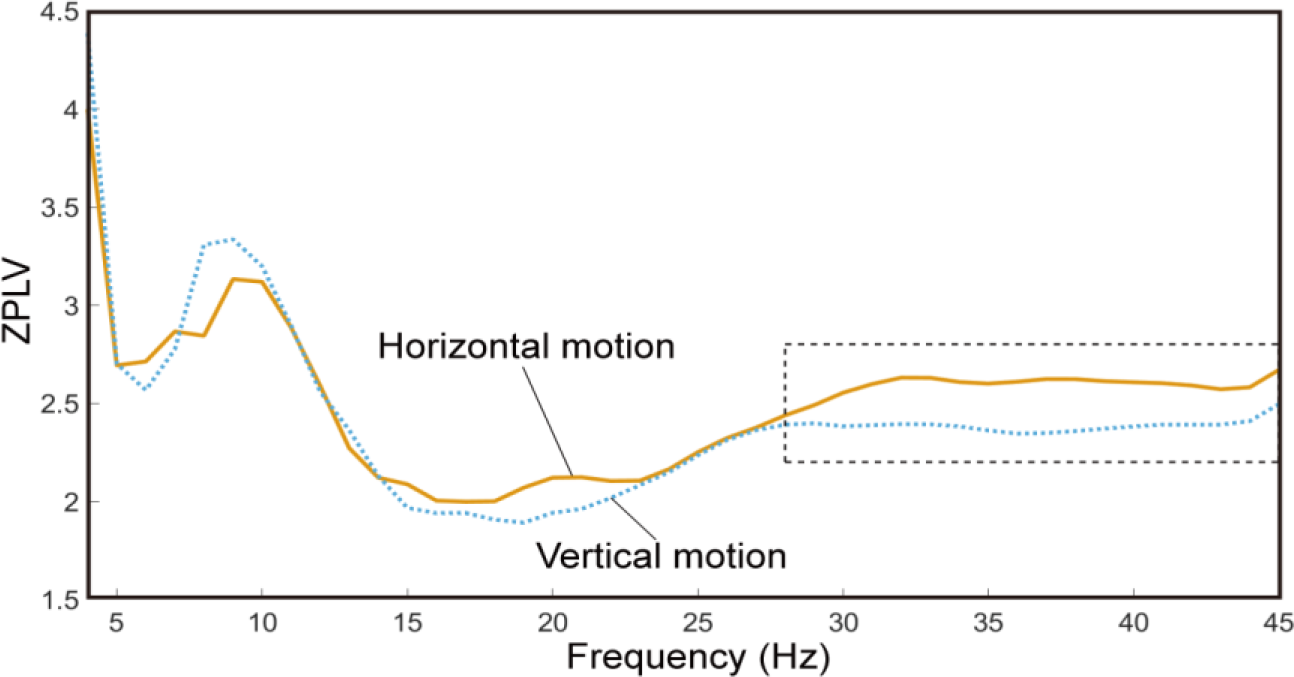
Interhemispheric phase synchrony in motion perception. The orange line indicates the group-averaged Rayleigh z value for the phase-locking value (*ZPLV*) during the perception of horizontal motion. The blue dashed line indicates the group-averaged *ZPLV* during the perception of vertical motion. *ZPLV* was averaged across the interhemispheric electrode pairs around hMT+ (P7-P8, P7-P6, P7-PO8, PO7-P8, P5-P6, PO7-P6, and PO7-PO8) and across participants. The black dashed line depicts a frequency window (28 Hz to 45 Hz) in which significant differences between *ZPLV* during horizontal motion and vertical motion were observed in at least one of the electrode pairs by a cluster-based permutation test (*P* = 0.020: one-tailed cluster test, *n* = 15, cluster-forming threshold *P* < 0.050, permutations = 10,000).

### Resting TMS-EEG for estimating intrinsic effective connectivity

We performed concurrent TMS-EEG recordings in the resting condition to estimate the intrinsic individual effective connectivity in oscillatory dynamics between left and right hemispheres. TMS pulses were delivered at intervals from 5 to 7 s with an intensity of 90% of the active motor threshold. Participants were given 90 TMS pulses for each target location (right hMT+, left hMT+, and sham: coil located 10 cm above the vertex) using an MRI-based TMS coil navigation system. We analyzed the effective connectivity between right hMT+ and left hMT+ by measuring the degree of interhemispheric propagation of frequency-specific TMS-induced phase resetting on an individual basis. TMS-induced phase resetting was estimated as the Rayleigh *z* value for the phase locking factor (*ZPLF).* We focused on the maximum *ZPLF* values between 0 ms and 100 ms after the TMS pulse in the ipsilateral hMT+ and the maximum *ZPLF* values between 15 ms and 150 ms in the contra-lateral hMT+ after the ipsilateral peak (Fig. 3). The *ZPLF* was averaged across the frequencies and electrodes corresponding to the gamma-band cluster that showed significant group difference between horizontal and vertical perception in the EEG (DDQ) experiment. The time courses of gamma-band *ZPLF* for hMT+ targeted TMS were statistically compared with those for the sham condition by a cluster-based permutation test (Maris and Oostenveld, 2007; Maris et al., 2007). We found significant increases in both contra- and ipsi-lateral *ZPLF* for the gamma band when TMS was delivered to the right hMT+ compared with the sham condition (P = 0.000 and *P* = 0.026: ipsilateral *ZPLF,* one-tailed cluster test, *n* = 12, 10,000 permutations, −39 ms to 81 ms and 89ms to 140 ms; *P* = 0.002: contralateral *ZPLF,* one-tailed cluster test, *n* = 12, 10,000 permutations, −11 ms to 84 ms, cluster-forming threshold *P* < 0.050: one-tailed *t*-test) (Fig. 3C). Group-averaged *ZPLF* differences showed peaks at 15 ms in the right hemisphere and at 66 ms in the left hemisphere. The latency of peak *ZPLF* differences averaged across participants was 23.3 ± 17.5 ms (mean ± s.d. *n* = 12) for the right hemisphere (ipsilateral to TMS), and 80.2 ± 28.0 ms (mean ± s.d. *n* = 12) for the left hemisphere (contralateral to TMS). We also observed significant increases in *ZPLF* in both contralateral and ipsilateral *ZPLF* in the gamma band when TMS was delivered to the left hMT+ compared with the sham condition (*P* = 0.0003: ipsilateral *ZPLF* one-tailed cluster test, *n* = 12, 10,000 permutations, −19 ms to 82 ms; *P* = 0.049 and *P* = 0.025: contralateral *ZPLF* one-tailed cluster test, *n* = 12, 10,000 permutations, 29 ms to 61 ms and 69 ms to 98 ms, cluster-forming threshold *P* < 0.050: one-tailed *t*-test) (Fig. 3D). Group-averaged *ZPLF* differences showed peaks at 14 ms in the left hemisphere and at 47 ms in the right hemisphere. The latency of peak *ZPLF* differences averaged across participants was 26.7 ± 23.7 ms (mean ± s.d. *n* = 12) for the left hemisphere (ipsilateral to TMS), and 78.3 ± 30.5 ms (mean ± s.d. *n* = 12) for the right hemisphere (contralateral to TMS). To assess interhemispheric effective connectivity in oscillatory dynamics, the normalized contralateral *ZPLF*_peak_ was calculated using the peak *ZPLF* values in both hemispheres.

**Figure 3.**
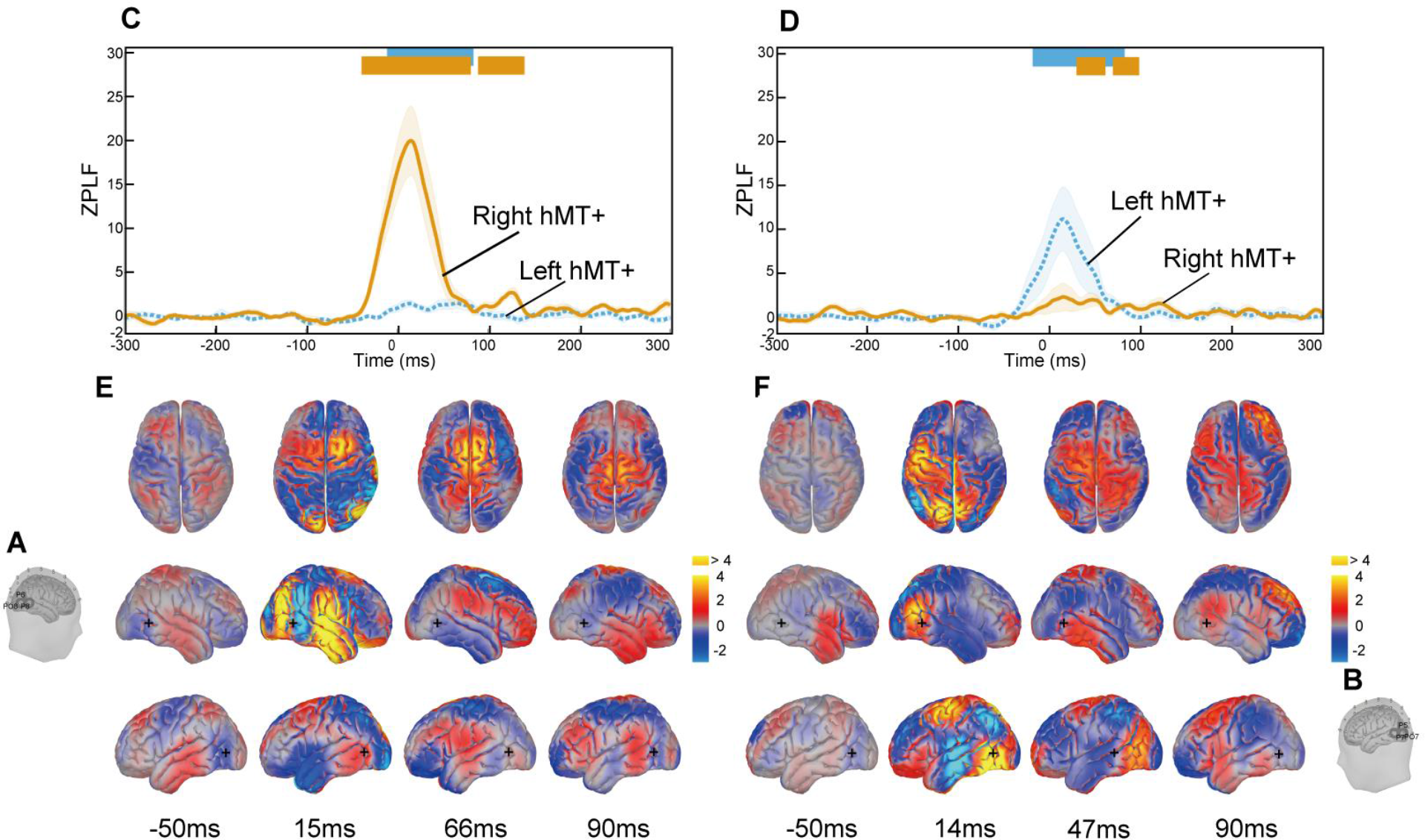
Global propagation of TMS-induced phase resetting. The TMS coil was located over right hMT+ (A) or left hMT+ (B). (**C**) and (**D**) Group-averaged (n = 12) time course of gamma-band *ZPLF* when TMS was delivered to the right hMT+ (**C**) and the left hMT+ (**D**). The shaded areas indicate the standard error of the mean (s.e.m.). The *ZPLF* time course was obtained by averaging *ZPLF* across the significant electrodes and frequencies corresponding to the significant cluster obtained in the EEG (DDQ) experiment after subtracting *ZPLF* in the sham condition from the real TMS condition. The thick blue lines at the top of the time-course plots indicate the times when the *ZPLF* for the left hMT+ was significantly larger during real TMS sessions than during sham sessions, as assessed by a cluster-based permutation test (*P* < 0.05: one-tailed cluster test, *n* = 12, 10,000 permutations; cluster-forming threshold: *P* < 0.05: one-tailed *t*-test). This is similar for the thick orange lines and the *ZPLF* for the right hMT+. (**E**) and (**F**) Corresponding topographic maps are shown below the time-course plots. Black crosses indicate the mean location of hMT+ in the sagittal plane.

### Linking effective connectivity, functional connectivity, and perception

We analyzed the correlations between perceptual biases and intrinsic effective connectivity in the gamma band and found a significant inter-individual positive correlation between the horizontal bias in DDQ perception and the gamma-band normalized contralatera*l ZPLF*_peak_ when TMS was delivered to right hMT+ (*P* = 0.012: one-tailed Spearman’s correlation,*ρ* = 0.713, *n* = 12, Bonferroni-corrected for 2 TMS-targeted sites) (Fig. 4A). In contrast to the right hMT+ targeted TMS, there was no significant correlation between perceptual bias and the gamma-band normalized contralateral *ZPLF*_peak_ when TMS was delivered to the left hMT+ (*P* = 0.366: one-tailed Spearman’s correlation *ρ* = 0.287, *n* = 12, Bonferroni-corrected for 2 TMS-targeted sites) (Fig. 4B). Moreover, we analyzed the correlation between perceptual bias and functional connectivity assessed as gamma-band phase synchrony (*ZPLV*) during inter-stimulus resting intervals using the same electrode pairs and frequency bands in the significant cluster. The *ZPLV* during rest was not significantly correlated with perceptual bias (*P* = 0.732: right hMT+ targeted TMS, one-tailed Spearman’s correlation *ρ* = −0.112, *P* = 0.766: left hMT+ targeted TMS, one-tailed Spearman’s correlation *n* = 12, *ρ* = 0.098, before Bonferroni-corrected for 2 TMS-targeted sites).

**Figure 4.**
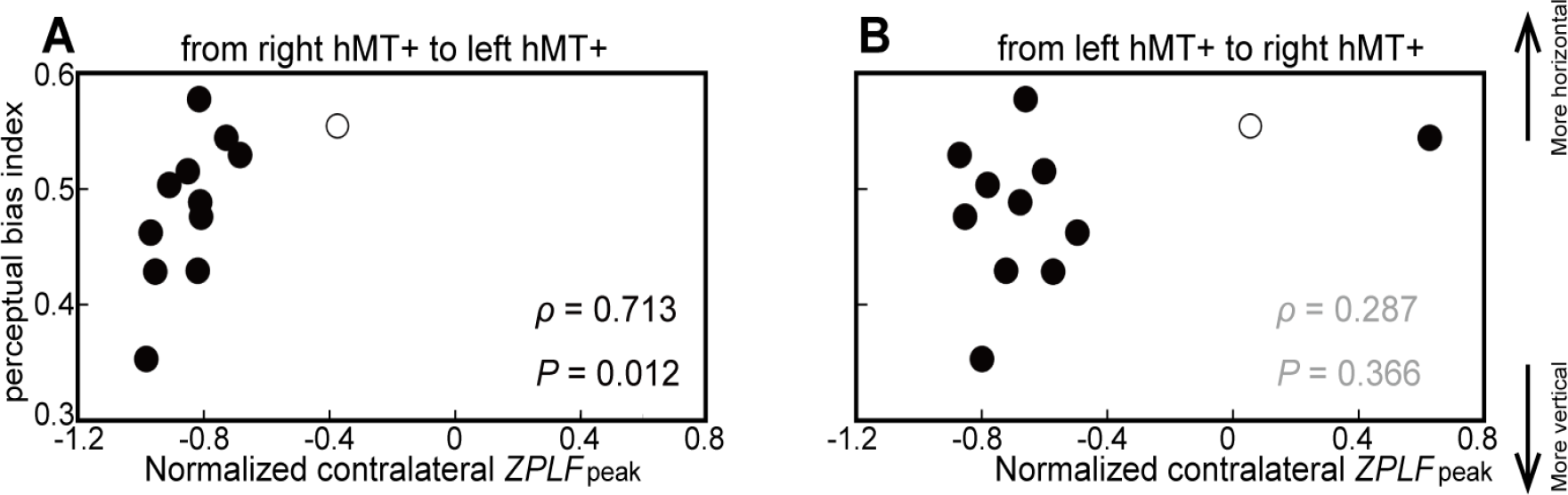
The relationship between individual perceptual bias and the contralateral propagation of phase resetting, estimated as normalized contralateral *ZPLF*_peak_. (**A**) The relationship between perceptual bias and the propagation from right hMT+ to left hMT+. A significant correlation between perceptual bias and the gamma-band normalized contralateral ZPLFpeak was observed (*P* = 0.012: one-tailed Spearman’s correlation, *ρ* = 0.713, *n* = 12, Bonferroni-corrected for 2 TMS-targeted sites) (**B**) The relationship between perceptual bias and propagation from the left hMT+ to the right hMT+. White circles indicate the individual who perceived a moving phosphene across the left and right visual fields. In contrast to the right hMT+ targeted TMS, there was no significant correlation between perceptual bias and the gamma-band normalized contralateral *ZPLF*_peak_ when TMS was delivered to the left hMT+ (*P* = 0.366: one-tailed Spearman’s correlation *ρ*= 0.287, *n* = 12, Bonferroni-corrected for 2 TMS-targeted sites)

Altogether, these results demonstrate that resting-state intrinsic effective connectivity assessed by the frequency-specific propagation of TMS-induced phase resetting is a robust methodology to predict individual behavioral bias in the DDQ perception task, which in turn is associated with the individual’s capacity to integrate information via the gamma band across left and right hMT+.

## Discussion

Here, we propose a novel technique to measure intrinsic effective connectivity and perceptual performance on an individual basis. Using a TMS-EEG co-registration methodology that utilizes individualized coil navigation and frequency-specific phase resetting, we demonstrate that individual human variation in perception can be tracked solely based on data from resting-state intrinsic effective connectivity in the dynamics of neural oscillations. Propagation of TMS-induced gamma-band phase resetting from right to left hMT+ explained individual bias in a perception task.

### TMS-EEG for probing effective connectivity in oscillatory dynamics

At the behavioral level, we found inter-individual correlations between horizontal motion bias and the propagation of phase resetting from the right to the left hMT+ in the gamma band. It is known that there are inter-individual differences in the ratio of the durations of horizontal and vertical motion perception (Chaudhuri and Glaser, 1991; Genç et al., 2011; Shimono et al., 2012). The results suggest that the underlying effective connectivity between oscillatory neural dynamics probed by our approach is causally responsible for individual differences in perception. However, it should be noted that functional connectivity of individual intrinsic gamma-band as measured by conventional resting-state gamma-band phase synchrony obtained using *zPLV* between electrodes over left and right hMT+ did not account for individual perceptual bias. The advantage of effective connectivity in our approach over current functional connectivity methods might be accounted for by a better signal-to-noise ratio of neural dynamics obtained by directly perturbing the relevant brain regions with greater precision. Moreover, probing effective connectivity by TMS-EEG manipulation can differentiate real and spurious communication between brain regions (Kitajo and Okazaki, 2016). We can only detect phase reset in a distant region when there is directional connectivity from the TMS-targeted region to the distant regions. In principle, if two regions are driven by a common neural oscillator without effective communication between them, we can detect functional connectivity (i.e. phase synchrony) but not effective connectivity. Therefore, our approach to estimate effective connectivity has a higher resolution than conventional functional connectivity methods for evaluating neural communication and predicting individual-level perception/cognition.

### Intrinsic effective connectivity and information integration

The correspondence we observed between intrinsic effective connectivity in brain dynamics probed by TMS-EEG during rest, and in the capacity to integrate information from perception suggests that effective connectivity of intrinsic network dynamics involved in frequency-specific spontaneous activity constrains the capacity to process information associated with perception. Large-scale phase synchronization of brain activity plays an important role in various perceptual and cognitive functions (Engel and Singer, 2001; Varela et al., 2001; Ward, 2003). Rose et al. found that gamma coherence between electrodes near the left and right hMT+ increased during perception of illusory horizontal motion in DDQ perception (Rose and Büchel, 2005). In marked contrast, we found enhanced interhemispheric phase synchrony between gamma-band CSD-transformed EEG signals over the right and left hMT+ during horizontal motion perception. It was reported that anti-phase transcranial alternating current stimulation (tACS) of both hemispheres in the gamma band modulates the duration of horizontal motion perception in DDQ (Helfrich et al., 2014). Horizontal motion perception is sustained longer by in-phase bilateral gamma-tACS and accompanied by increased interhemispheric EEG coherence in the gamma-band while vertical motion perception is reinforced by decreased interhemispheric coherence in the same gamma-band (Helfrich et al., 2014).

These prior studies focused on task-related neural communication in DDQ perception, and support our results showing that participants who exhibited greater gamma-band effective connectivity from the right to the left hMT+ probed by TMS-EEG at rest showed a greater capacity for integration of information across hemispheres, as illustrated by the greater degree of horizontal bias in DDQ perception. The results further indicate that intrinsic effective connectivity in gamma-band oscillatory dynamics between two separate areas (i.e., left and right hMT+) at rest is the actual mechanistic basis shaping the participant’s perception, in which information from the two areas needs to be integrated because visual hemifields are segregated in right and left visual cortices.

We also found an interesting laterality effect, where the propagation of phase resetting from right hMT+ to left hMT+ explained horizontal DDQ motion perception, but propagation of phase resetting from left hMT+ to right hMT+ did not. This result may be related to the differences in large-scale connectivity and information integration between the two hemispheres. Gotts et al. used fMRI resting-state functional connectivity analyses to show that the left hemisphere interacts more within the same hemisphere, whereas the right hemisphere interacts with both hemispheres suggesting that the right hemisphere is more integrative while the left is more segregated (Gotts et al., 2013).

### Strengths and limitations of TMS-EEG approaches in behavioral studies

We believe that the TMS-EEG manipulative technique we describe here will allow direct access to a wide variety of behavioral measures that require resting-state effective connectivity for tracking individual differences in perception or cognition via the long-range integration of information. In this regard, our method will allow TMS-EEG to complement other imaging methods like high resolution fMRI combined with decoding in dissecting individual variation. Moreover, individual-level effective connectivity methods will have major clinical implications for personalized medicine in diseases that are known to be associated with abnormal or impaired neural synchrony, including epilepsy, schizophrenia (Uhlhaas et al., 2006), autism (Murias et al., 2007), and stroke (Kawano et al., 2017).

The methodology we describe also has several limitations at present. The precise anatomical pathways for the propagation of phase resetting information remain unclear, as bilateral hMT+ areas are connected by several pathways and the corpus callosum (CC) mediates connectivity between the left and right MT regions in monkeys (Maunsell and van Essen, 1983). A diffusion tensor imaging (DTI) study showed that inter-individual variability in DDQ perception can be partially accounted for by the microstructural properties of fibers in the posterior CC linking left and right hMT+ (Genç et al., 2011). In addition, interhemispheric EEG coherence in the gamma-band is mediated by the CC (Kiper et al., 1999). Subcortically, the superior colliculus (SC) and the pulvinar nucleus (PN) are associated with the connection between the left and right hMT+ in DDQ perception (Shimono et al., 2012) and hMT+ is also directly connected with the lateral geniculate nucleus (LGN) of the thalamus (Gaglianese et al., 2012). Whether the TMS-induced phase resetting from the ipsilateral to contralateral hMT+ propagates through the CC, SC, PN, and LGN remains to be investigated using multi-modal imaging to complement TMS-EEG. For instance, effective connectivity probed by TMS-EEG and it’s concordance with the degree of anatomical connections by DTI as well as functional connectivity using fMRI may help to reveal the underlying neural connections.

TMS-EEG also has limited spatial resolution and target location at present. It is possible to target TMS to bilateral hMT+ because it is located in superficial areas of the cortex. Obviously, it will be more difficult to target cortical areas which are farther from the scalp surface (Siebner et al., 2010). Therefore, the types of behavioral tasks will be limited with respect to target locations. For example, it will be hard to directly target the fusiform face area, which is located on the ventral surface of the temporal lobe, to investigate effective connectivity associated with face processing. State-dependent components can also be confounding factors in these studies. It is well known that TMS-evoked responses show brain-state-dependent changes in sleep and wake states (Massimini et al., 2005). Although our participants were awake, changes in arousal level can affect the results.

Despite these issues, we believe that our individualized co-registration technique will enable a new generation of TMS-EEG studies that will focus on quantitative individual participant behavioral dissection via direct manipulation of nonlinear neural dynamics mediating perceptual, motor, and cognitive processes. Our perturbation method will enhance the experimental toolbox for quantitative brain physiology and behavior needed for deeper mechanistic insight into complex individual patterns of variation in human behavior. Furthermore, combining our TMS-EEG co-registration technique with a closed-loop EEG feedback system (Zrenner et al., 2016) for dynamics-dependent stimulation of specific brain areas will enable the manipulative dissection of neural dynamics and brain functions in real-time and neurofeedback experimental or therapeutic configurations.

## Acknowledgements

Y.M. was supported by a Grant-in-Aid for JSPS Fellows (26-8352), the RIKEN Junior Research Associate (JRA) program. K.K. was supported by a research grant from the Toyota Motor Corporation, JST PRESTO, and MEXT KAKENHI Grant Number JP15H05877. We thank Yutaka Uno, and Toshihisa Tanaka for advice and comments, and Jumpei Mori and Tatsuhiko Sone for help with the data analysis. We thank Dr. Charles Yokoyama at RIKEN Brain Science Institute for his critical comments and editing of the manuscript.

## References

Bortoletto M, Veniero D, Thut G, Miniussi C (2015) The contribution of TMS-EEG coregistration in the exploration of the human cortical connectome. Neurosci Biobehav Rev 49:114–124.

Bullmore E, Sporns O (2009) Complex brain networks: graph theoretical analysis of structural and functional systems. Nat Rev Neurosci 10:186–198.

Chaudhuri A, Glaser DA (1991) Metastable motion anisotropy. Vis Neurosci 7:397–407.

Dumoulin SO, Bittar RG, Kabani NJ, Baker CLJ, Le Goualher G, Bruce Pike G, Evans AC (2000) A new anatomical landmark for reliable identification of human area V5/MT: a quantitative analysis of sulcal patterning. Cereb Cortex 10:454–463.

Engel AK, Singer W (2001) Temporal binding and the neural correlates of sensory awareness. Trends Cogn Sci 5:16–25.

Friston KJ (2011) Functional and effective connectivity: a review. Brain Connect 1:13–36.

Gaglianese A, Costagli M, Bernardi G, Ricciardi E, Pietrini P (2012) Evidence of a direct influence between the thalamus and hMT+ independent of V1 in the human brain as measured by fMRI. NeuroImage 60:1440–1447.

Genç E, Bergmann J, Singer W, Kohler A (2011) Interhemispheric connections shape subjective experience of bistable motion. Curr Biol 21:1494–1499.

Gotts SJ, Jo HJ, Wallace GL, Saad ZS, Cox RW, Martin A (2013) Two distinct forms of functional lateralization in the human brain. Proc Natl Acad Sci U S A 110:3435–3444.

Guzman-Lopez J, Silvanto J, Seemungal BM (2011) Visual motion adaptation increases the susceptibility of area V5/MT to phosphene induction by transcranial magnetic stimulation. Clin Neurophysiol 122:1951–1955.

Helfrich RF, Knepper H, Nolte G, Strüber D, Rach S, Herrmann CS, Schneider TR, Engel AK (2014) Selective modulation of interhemispheric functional connectivity by HD-tACS shapes perception. PLoS Biol 12:e1002031.

Herring JD, Thut G, Jensen O, Bergmann TO (2015) Attention modulates TMS-locked alpha oscillations in the visual cortex. J Neurosci 35:14435–14447.

Hu X, Le TH, Parish T, Erhard P (1995) Retrospective estimation and correction of physiological fluctuation in functional MRI. Magn Reson Med 34:201–212.

Ilmoniemi RJ, Virtanen J, Ruohonen J, Karhu J, Aronen HJ, Näätänen R, Katila T (1997) Neuronal responses to magnetic stimulation reveal cortical reactivity and connectivity. Neuroreport 8:3537–3540.

Kawano T, Hattori N, Uno Y, Kitajo K, Hatakenaka M, Yagura H, Fujimoto H, Yoshioka T, Nagasako M, Otomune H, Miyai I (2017) Large-scale phase synchrony reflects clinical status after stroke: An EEG study. Neurorehabil Neural Repair 31:561–570.

Kawasaki M, Uno Y, Mori J, Kobata K, Kitajo K (2014) Transcranial magnetic stimulation-induced global propagation of transient phase resetting associated with directional information flow. Front Hum Neurosci 8:173.

Kayser J, Tenke CE (2006) Principal components analysis of Laplacian waveforms as a generic method for identifying ERP generator patterns: I. Evaluation with auditory oddball tasks. Clin Neurophysiol 117:348–368.

Kiper DC, Knyazeva MG, Tettoni L, Innocenti GM (1999) Visual stimulus-dependent changes in interhemispheric EEG coherence in ferrets. J Neurophysiol 82:3082–3094.

Kitajo K, Hanakawa T, Ilmoniemi RJ, Miniussi C (2015) A contemporary research topic: manipulative approaches to human brain dynamics. Front Hum Neurosci 9.

Kitajo K, Okazaki Y (2016) TMS-EEG for probing distinct modes of neural dynamics in the human brain. Adv Cogn Neurodyn (V):211–216.

Koessler L, Maillard L, Benhadid A, Vignal JP, Felblinger J, Vespignani H, Braun M (2009) Automated cortical projection of EEG sensors: anatomical correlation via the international 10-10 system. NeuroImage 46:64–72.

Komssi S, Aronen HJ, Huttunen J, Kesäniemi M, Soinne L, Nikouline V V., Ollikainen M, Roine RO, Karhu J, Savolainen S, Ilmoniemi RJ (2002) Ipsi- and contralateral EEG reactions to transcranial magnetic stimulation. Clin Neurophysiol 113:175–184.

Lachaux JP Rodriguez E, Martinerie J, Varela FJ (1999) Measuring phase synchrony in brain signals. Hum Brain Mapp 8:194–208.

Maris E, Oostenveld R (2007) Nonparametric statistical testing of EEG- and MEG-data. J Neurosci Methods 164:177–190.

Maris E, Schoffelen JM, Fries P (2007) Nonparametric statistical testing of coherence differences. J Neurosci Methods 163:161–175.

Massimini M, Ferrarelli F, Huber R, Esser SK, Singh H, Tononi G (2005) Breakdown of cortical effective connectivity during sleep. Science 309:2228–2232.

Maunsell JH, van Essen DC (1983) The connections of the middle temporal visual area (MT) and their relationship to a cortical hierarchy in the macaque monkey. J Neurosci 3:2563–2586.

Mognon A, Jovicich J, Bruzzone L, Buiatti M (2010) ADJUST: An automatic EEG artifact detector based on the joint use of spatial and temporal features. Psychophysiology 48:229–240.

Morishima Y, Akaishi R, Yamada Y, Okuda J, Toma K, Sakai K (2009) Task-specific signal transmission from prefrontal cortex in visual selective attention. Nat Neurosci 12:85–91.

Murias M, Swanson JM, Srinivasan R (2007) Functional connectivity of frontal cortex in healthy and ADHD children reflected in EEG coherence. Cereb Cortex 17:1788–1799.

Nikouline V, Ruohonen J, Ilmoniemi RJ (1999) The role of the coil click in TMS assessed with simultaneous EEG. Clin Neurophysiol 110:1325–1328.

Oostenveld R, Fries P, Maris E, Schoffelen JM (2011) FieldTrip: Open source software for advanced analysis of MEG, EEG, and invasive electrophysiological data. Comput Intell Neurosci:156869.

Perrin F, Pernier J, Bertrand O, Echallier JF (1989) Spherical splines for scalp potential and current density mapping. Electroencephalogr Clin Neurophysiol 72:184–187.

Perrin F, Pernier J, Bertrand O, Echallier JF (1990) Corrigenda EEG 02274. Electroencephalogr Clin Neurophysiol 76:565–565.

Rodriguez E, George N, Lachaux JP, Martinerie J, Renault B, Varela FJ (1999) Perception’s shadow: long-distance synchronization of human brain activity. Nature 397:430–433.

Rosanova M, Casali A, Bellina V, Resta F, Mariotti M, Massimini M (2009) Natural frequencies of human corticothalamic circuits. J Neurosci 29:7679–7685.

Rose M, Büchel C (2005) Neural coupling binds visual tokens to moving stimuli. J Neurosci 25:10101–10104.

Ruzzoli M, Marzi C a, Miniussi C (2010) The neural mechanisms of the effects of transcranial magnetic stimulation on perception. J Neurophysiol 103:2982–2989.

Shimono M, Kitajo K, Takeda T (2011) Neural processes for intentional control of perceptual switching: a magnetoencephalography study. Hum Brain Mapp 32:397–412.

Shimono M, Mano H, Niki K (2012) The brain structural hub of interhemispheric information integration for visual motion perception. Cereb Cortex 22:337–344.

Siebner HR, Hartwigsen G, Kassuba T, Rothwell J (2010) How does transcranial magnetic stimulation modify neuronal activity in the brain? - Implications for studies of cognition. Cortex 45:1035–1042.

Sterzer P, Eger E, Kleinschmidt A (2003) Responses of extrastriate cortex to switching perception of ambiguous visual motion stimuli. Neuroreport 14:2337–2341.

Tadel F, Baillet S, Mosher JC, Pantazis D, Leahy RM (2011) Brainstorm: a user-friendly application for MEG/EEG analysis. Comput Intell Neurosci 2011:879716.

Tallon-Baudry C, Bertrand O, Delpuech C, Pernier J (1996) Stimulus specificity of phase-locked and non-phase-locked 40 Hz visual responses in human. J Neurosci 16:4240–4249.

Thut G, Veniero D, Romei V, Miniussi C, Schyns P, Gross J (2011) Rhythmic TMS causes local entrainment of natural oscillatory signatures. Curr Biol 21:1176–1185.

Uhlhaas PJ, Linden DEJ, Singer W, Haenschel C, Lindner M, Maurer K, Rodriguez E (2006) Dysfunctional long-range coordination of neural activity during gestalt perception in Schizophrenia. J Neurosci 26:8168–8175.

Varela F, Lachaux JP, Rodriguez E, Martinerie J (2001) The brainweb: Phase synchronization and large-scale integration. Nat Rev Neurosci 2:229–239.

Ward LM (2003) Synchronous neural oscillations and cognitive processes. Trends in Cognitive Sciences 7:553–559.

Zrenner C, Belardinelli P, Müller-Dahlhaus F, Ziemann U (2016) Closed-loop neuroscience and non-invasive brain stimulation: A tale of two loops. Front Cell Neurosci 10:1–7.

